# Exogenous L-lactate promotes astrocyte plasticity but is not sufficient for enhancing striatal synaptogenesis or motor behavior in mice

**DOI:** 10.1101/2020.04.13.039446

**Authors:** Adam J. Lundquist, Tyler J. Gallagher, Giselle M. Petzinger, Michael W. Jakowec

## Abstract

L-lactate is an energetic and signaling molecule that is key to the metabolic and neuroplastic connection between astrocytes and neurons and may be involved in exercise-induced neuroplasticity. This study sought to explore the role of L-lactate in astrocyte reactivity and neuroplasticity. Using in vitro cultures of primary astrocytes, we show L-lactate increased expression of plasticity-related genes, including neurotrophic factors, *Bdnf, Gdnf, Cntf* and the immediate early gene *cFos*. L-lactate’s promotion of neurotrophic factor expression may be mediated in part by the lactate receptor HCAR1 since application of the HCAR1 agonist 3,5-DHBA also increased expression of *Bdnf* in primary astrocytes. In vivo L-lactate administration to healthy mice caused a similar increase in the expression of plasticity-related genes as well as increased astrocyte morphological complexity in a region-specific manner, with increased astrocytic response found in the striatum but not the ectorhinal cortex, regions of the brain where increases in regional cerebral blood flow are increased or unaltered, respectively, with motor behavior. Additionally, L-lactate administration did not cause synaptogenesis or improve motor behavior based on the latency to fall on the accelerating rotarod, suggesting that L-lactate administration can initiate astrocyte-specific gene expression, but the activation of motor circuits is necessary to initiate striatal neuroplasticity. These results suggest that peripheral L-lactate is likely an important molecular component of exercise-induced neuroplasticity by acting in an astrocyte-specific manner to prime the brain for neuroplasticity.

## Introduction

Peripheral sources of L-lactate, resulting from glucose metabolism in muscle, are known to play a critical role in the central nervous system (CNS). For example, L-lactate administration in mice recapitulates aspects of intensive aerobic exercise and has brain- and liver-specific effects on metabolism (E, Lu, Selfridge, Burns, & Swerdlow, 2013). Additionally, L-lactate administration has antidepressant effects in mice, similar to aerobic exercise training (Carrard et al., 2018; Duman, Schlesinger, Russell, & Duman, 2008). Intensive aerobic exercise causes a significant increase in L-lactate concentrations in blood plasma; exercise also increases regional cerebral blood flow (Wang et al., 2013), and allows L-lactate to act as a metabolic substrate for the brain during exercise (Ide, Schmalbruch, Quistorff, Horn, & Secher, 2000; van Hall et al., 2009). Thus, L-lactate production in muscles and L-lactate metabolism in the brain operate in a cooperative fashion linking body activity and brain bioenergetics during motor behavior. While the impact of physiologic levels of L-lactate (∼10mM, Morland et al., 2017) on neuronal function and signaling is well-known (Yang et al., 2014),the relative impact of L-lactate on astrocytes is less well understood.

Within the CNS, L-lactate is preferentially produced by astrocytes through aerobic glycolysis (Magistretti & Allaman, 2018), and has been shown to be a critical component for memory formation and plasticity by acting in an energetic substrate for neurons through the astrocyte-neuron lactate shuttle (ANLS) (Suzuki et al., 2011). L-lactate can also act as a signaling molecule through its activation of the Gi/o-protein coupled receptor hydroxycarboxylic acid receptor 1 (HCAR1) promoting angiogenesis in vivo (Bergersen, 2015; Morland et al., 2017) and regulating neuronal excitability (Abrantes et al., 2019). Astrocytic endfeet tile the entirety of brain vasculature, where they regulate blood flow and substrate uptake through the blood brain barrier. Both astrocytes and brain endothelial cells express monocarboxylate transporters (MCTs) capable of transporting L-lactate from the peripheral blood stream into the brain (Lauritzen et al., 2014), suggesting that astrocytes regulate the uptake of muscle-derived L-lactate. However, how physiologic L-lactate influences astrocytic function and astrocyte-specific gene expression in neuroplasticity remains a major gap in knowledge.

The purpose of this study was to investigate the effects of physiologic L-lactate on astrocyte-specific gene expression and morphology in both astrocyte cell cultures and in normal healthy mice. Studies were focused on two brain regions, the striatum (STR) and ectorhinal cortex (ETC) since previous studies showed exercise-induced increase (STR) or no change (ETC) in regional cerebral blood flow, respectively (Wang et al., 2013). We also investigated whether L-lactate could promote synaptogenesis in these regions and improve motor performance. These findings reveal potential underlying mechanisms by which peripheral L-lactate may impact the CNS acting in part through astrocytes to mediate neuroplasticity.

## Materials and Methods

### Animals

No pre-registration was performed for this study. C57BL/6J mice 10 to 14-weeks of age (The Jackson Laboratory, Bar Harbor, ME) were used for this study and were housed at the University of Southern California vivarium. Mice were housed in groups of 4 to 5 per cage and maintained on a reverse 12-hour light/dark cycle (lights off at 0700 hours) with ad libitum access to food and water. Mice of both sexes were used in equal proportions. Postnatal day 0-4 (PND0-4) pups from in-house C57BL/6J breeding pairs were used for the generation of primary astrocyte cultures, which are described in more detail below. All procedures were approved by the Institutional Animal Care and Use Committee of the University of Southern California (IACUC protocol #9766, #21044) and conducted in accordance with the National Research Council’s Guide for the Care and Use of Laboratory Animals (Committee for the Update of the Guide for the Care and Use of Laboratory Animals; National Research Council, 2010). No randomization was explicitly performed to allocate subjects in the study.

### Experimental Design and In Vivo Lactate Administration

An overview of experimental design and relevant details are included in Figure 1, including planned endpoints for all groups. All experiments were conducted between 0900 and 1400 hours. Mice (n = 32) were divided into two groups for all experiments (n = 16 per group) and intraperitoneal (I.P.) injected daily for 10 days with either a solution of sodium L-lactate (Cat# 71718, Millipore-Sigma, St. Louis, MO; 2g/kg body weight, estimated final concentration 10mM) (Morland et. al, 2017), dissolved in 0.9% saline, or an equal volume of vehicle (0.9% saline). After each daily injection, mice were placed on motorized treadmills (EXER-6M, Columbus Instruments, Columbus, OH) to walk for 10 min/day at a speed of 6m/min. Mice that received vehicle injections were placed on an adjacent treadmill to walk for 10 min/day at a speed of 6m/min. All mice were maintained at this low level of activity on the treadmill where L-lactate levels have been reported to not increase in rodents (Billat, Mouisel, Roblot, & Melki, 2005; Soya et al., 2007). All mice were monitored and handled daily and continued to be included in the study if they exhibited normal behavior and were otherwise healthy. No mice required veterinary care, died, or had to be otherwise excluded during the course of experiments.

**Figure 1.**
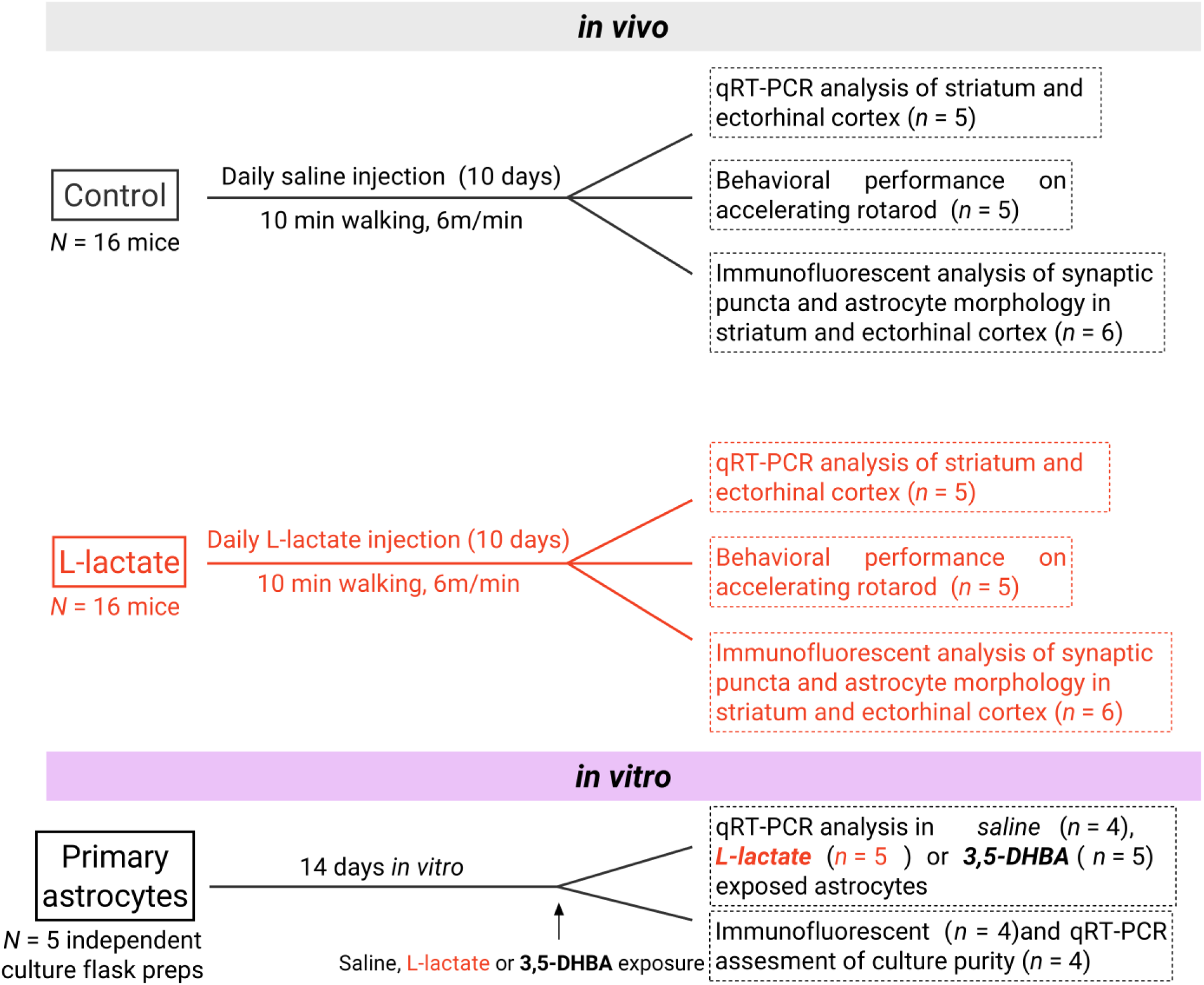
Overview of experimental design.

### Rotarod Motor Performance

Motor performance following in vivo L-lactate administration was tested on an accelerated rotarod using a modification of previously published methods (Rothwell et al., 2014). Following L-lactate administration as detailed above, a subset of mice (n = 5 per group) were trained on an accelerating rotarod (3cm diameter rod, divided into five lanes; Ugo Basile, Comerio, Italy). The rotarod accelerated over the course of 300 seconds from 6 to 60 rpm, and speed at time of fall and latency to fall were automatically recorded by magnetic trip plates. Mice were acclimated to the rotarod for 90 seconds before the start of their first trial of each day and were trained for five trials per day for four days, with a one-minute intertrial interval. A trial ended when the mouse made a complete backward revolution, fell off, or reached the 300 second threshold. Rotarod training was conducted by persons blind to the experimental group assignment of the mice.

### Brain Tissue Collection

Whole brains were collected on the last day of injection and walking. After the cessation of walking, mice were divided into groups for either fresh tissue collection (n = 6 mice, see below) or transcardial perfusion (n = 5 mice). For transcardial perfusion, one hour after the final session, mice were heavily sedated with intraperitoneal injections of tribromoethanol (250 mg/kg body weight) and assessed for lack of toe-pinch response. The use of tribromoethanol for mouse sedation is standard and approved for use in our protocol. Mice were transcardially perfused with 50 ml of ice-cold 0.9% saline followed by 100 ml of ice-cold 4% paraformaldehyde in phosphate buffered saline, pH 7.2 (PFA-PBS). Brains were extracted, transferred to 4% PFA-PBS at 4°C overnight and then to a 20% sucrose cryoprotection solution. After sinking, brains were rapidly frozen in 2-methylbutane cooled on dry ice and stored at −80°C until use.

Fresh tissue dissections were carried out immediately following the final session of injection and walking. Mice from both vehicle and L-lactate groups (n = 6 mice each) were sacrificed by cervical dislocation, as approved in our protocol and to limit potential effects of anesthesia on gene expression, and the brains were resected to ice-cold PBS, pH 7.2. Brains were microdissected to isolate the striatum (STR; Bregma +1.2 to −0.2mm A.P., including tissue bordered ventrally by the anterior commissure, dorsally by the corpus callosum, medially by the lateral ventricle, and ±2.5 mm laterally from the midline) (Halliday, Abeydeera, Lundquist, Petzinger, & Jakowec, 2019; Kintz et al., 2013) and ectorhinal cortex (ETC; Bregma −2.0 to −3.0 mm A.P., laterally bounded by the edge of the cortex and extending 1mm medially to the edge of the striatum, and −3.0 to −3.5 mm D.V.) (Halliday et al., 2019).

### Primary Astrocyte Isolation and Culture

Mouse astrocytes were prepared as previously described (Jouroukhin et al., 2018). Briefly, whole brains were removed from PND0-4 mice to ice-cold Dulbecco’s phosphate buffered saline (DPBS) and the cortices were freed of meninges, microdissected, and minced into small chunks. Cortices were digested with 0.25% trypsin-EDTA (Cat.# 25-510, Genesee Scientific, San Diego, CA) in DPBS at 37°C for 20 minutes with inversion every 5 minutes. Tissue lysate was centrifuged at 300 x g for 5 min, supernatant removed, and tissue pellet resuspended in astrocyte media (DMEM with 10% fetal bovine serum and 1% penicillin/streptomycin; Cat.# 25-500, Cat.# 25-550, Cat.# 25-512, Genesee Scientific). Tissue pellet was triturated with decreasing size pipettes until a single cell suspension was achieved, which was passed through a 40μm filter to strain large clumps of debris. The resulting single cell suspension was plated on 75 cm2 tissue-culture treated flasks and grown in a 37°C incubator with 5% CO2. Loosely attached glial cells were washed with DPS and removed by vigorously shaking flasks by hand for 2 minutes before exchanging DPBS with fresh astrocyte media. Astrocytes were confluent by DIV7, when cultures were incubated with 0.25% trypsin-EDTA at 37°C and plated on poly-D-lysine (PDL) coated coverslips or 6- or 12-well tissue-culture treated plates until future analysis. Astrocytes were confluent and used for experimentation by DIV14±1 day. Resultant astrocyte cultures were highly pure (on average >96% astrocytes) and purity was assessed through SOX9, GFAP, and ALDH1L1 immunocytochemistry and quantitative real-time PCR of cell-type specific genes as described below (Figure 2).

**Figure 2.**
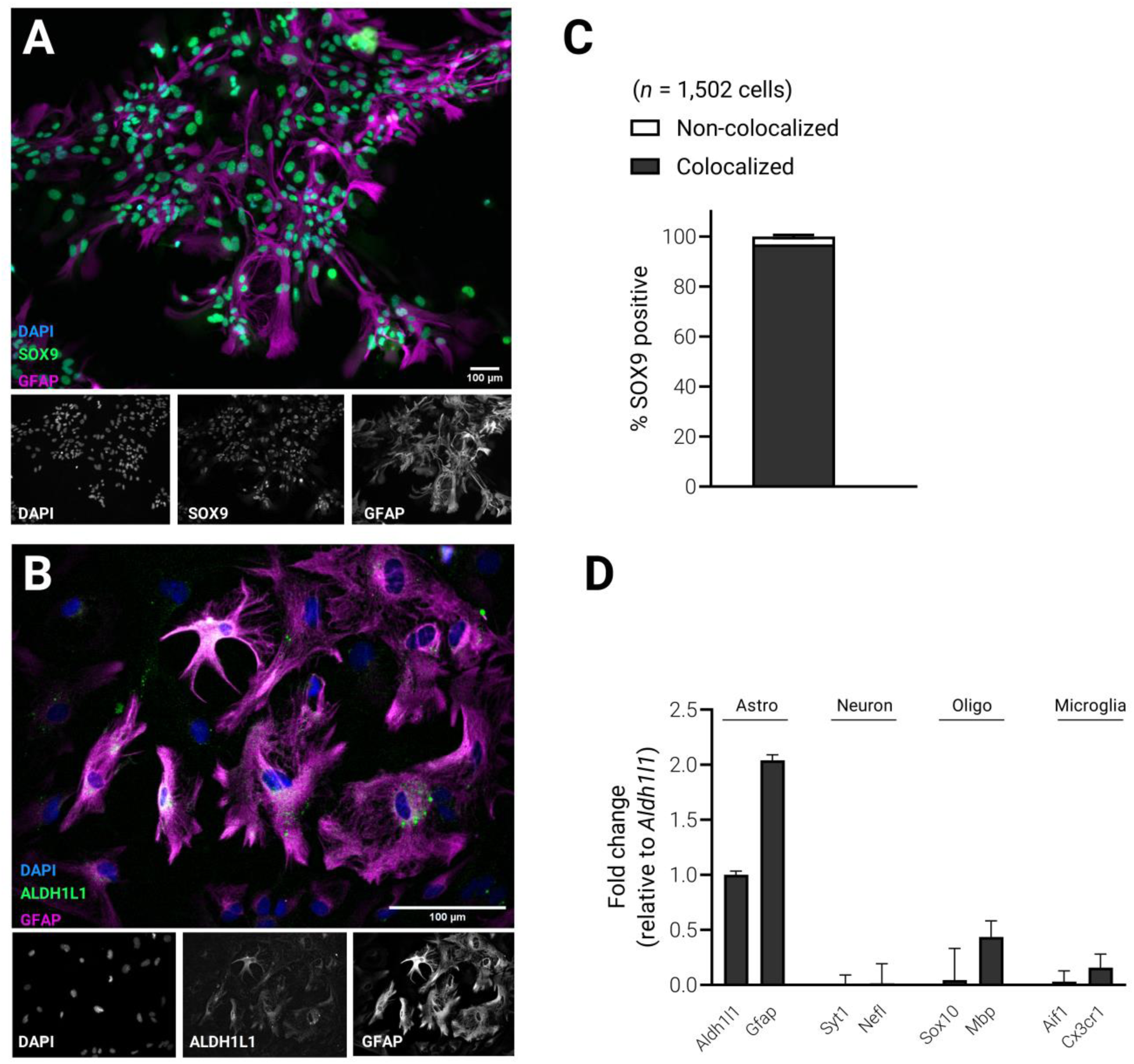
Primary mouse astrocyte cultures are highly pure. (A-B) Representative immunocytochemistry of primary astrocyte cultures demonstrate characteristic nuclear and cytoplasmic markers of astrocytes. (C) Over 96% of all DAPI-stained nuclei colocalized with the astrocytic nuclear marker SOX9 (*n* = 10 coverslips from four independent culture experiments). (D) Cell-specific transcriptional analysis with qRT-PCR verified astrocyte-enriched transcripts with very little non-astrocyte lineage gene expression (*n* = 4 samples from four independent culture experiments).

### In vitro administration of compounds

All experimental compounds were prepared in 50 or 100x stock solutions in 0.9% saline and kept frozen at −20°C until the day of experimentation. Primary astrocytes in 6- or 12-well plates had media exchanged on the day of experimentation and then stock solutions were added directly to culture media to their final concentration for 1 hour (unless otherwise noted). Final concentration for L-lactate was 10mM dissolved in 0.9% saline, while the final concentration of 3,5-DHBA (3,5-dihydroxybenzoic acid) was 2.5mM dissolved in 0.9% saline.

### Immunofluorescent Staining

#### Immunohistochemical Staining

Whole brains (fixed) were sliced in coronal sections (30 μm thickness) on a sliding freezing microtome (Leica CM1900, Leica Microsystems, Wetzlar, Germany) as previously described (Lundquist, Parizher, Petzinger, & Jakowec, 2019). Briefly, sections were washed in Tris-buffered saline, pH 7.2, with 0.2% Triton X-100 (TBST), blocked in 4% normal goat serum (NGS; Vector Laboratories Cat# S-1000, RRID:AB_2336615) in TBST, and incubated overnight at 4°C in primary antibody (2% NGS in TBST) with either rabbit anti-GFAP (1:2000, Agilent Cat# Z0334, RRID: AB_10013382), rabbit anti-synaptophysin (IgG, 1:2000, Abcam Cat# ab32127, RRID:AB_2286949) or mouse anti-PSD-95 (IgG1, 1:2000, Millipore Cat# AB9708, RRID:AB_2092543). Sections were washed with TBST and incubated in secondary antibody (2% NGS in TBST) with Alexa 568-conjugated goat anti-rabbit (1:5000, Thermo Fisher Scientific, Cat# A-11011, RRID: AB_143157), or Alexa 568-conjugated goat anti-mouse IgG1 (1:5000, Thermo Fischer Scientific, Cat# A-21124, RRID: AB_2535766). Sections were washed, mounted onto gelatin-coated slides, and coverslipped (Vectashield Hardset Antifade with DAPI; H-1500, Vector Labs, Burlingame, CA). Confocal images were taken on an IXB-DSU spinning disk Olympus BX-61 (Olympus America, Melville, NY) and captured with an ORCA-R2 digital CCD camera (Hamamatsu, Bridgewater, NJ) and MetaMorph Advanced software (Molecular Devices, San Jose, CA).

#### Immunocytochemical Staining

Primary astrocytes grown on PDL-coated coverslips were washed twice with ice-cold DPBS before being fixed with ice-cold 4% PFA-PBS (pH 7.2) for 10 minutes. Astrocytes were washed with TBS, permeabilized with TBST, and washed again with TBS. Astrocytes were blocked with 10% normal goat serum (NGS; S-1000, Vector Labs, Burlingame, CA) in TBST before incubating overnight at 4°C in primary antibody solution (4% NGS in TBST) with rat anti-GFAP (Millipore Cat# 345860-100UG, RRID:AB_211868), rabbit anti-GFAP (1:2000, Agilent Cat# Z0334, RRID: AB_10013382), rabbit anti-SOX9 (1:2000, Millipore Cat# AB5535, RRID:AB_2239761) and mouse anti-ALDH1L1 (1:500, UCDavis/NIH NeuroMab Facility Cat# 75-164, RRID:AB_10671695). Astrocytes were washed the following day with TBS and incubated in secondary antibody solution (2% NGS in TBST) with goat anti-mouse 488 (1:5000, Thermo Fisher Scientific Cat# A-11001, RRID:AB_2534069), goat anti-rabbit 488 (1:5000, Thermo Fisher Scientific Cat# A-11008, RRID:AB_143165), goat anti-rat 568 (1:5000, Thermo Fisher Scientific Cat# A-11077, RRID:AB_2534121) or goat anti-rabbit 568 (1:5000, Thermo Fisher Scientific, Cat# A-11011, RRID: AB_143157).

### Analyzing Synaptic Puncta

Synaptophysin- and PSD-95-positive puncta were captured in the striatum and ectorhinal cortex with a 40x objective and identical camera settings, and images were analyzed using the following workflow. First the background was subtracted, then images were manually thresholded at approximately the upper 1% of signal to eliminate non-specific or overlapping puncta. Finally, the Analyze Particles plugin for ImageJ (Schindelin et al., 2012) was used to quantify puncta for the entire field of view of the image. Two or more tissue sections through each anatomical region were analyzed per animal (n = 6 mice per group). Synaptic puncta were analyzed by persons blind to the experimental group.

### SOX9 Colocalization for Culture Purity

Assessment of relative purity of primary astrocytes was performed through colocalization analysis of SOX9-DAPI double positive nuclei according to published methods (Manders, Verbeek, & Aten, 1993). Briefly, two to three coverslips from four separate experiments with astrocytes stained with SOX9 and DAPI were captured with a 20x objective before using the Colocalization plugin for ImageJ for relative percentage of green/blue signal colocalization.

### Sholl Analysis

Morphological analysis of GFAP-positive astrocytes in the striatum and ectorhinal cortex were assessed using Sholl analysis as previously described (Lundquist et al., 2019; Sholl, 1953). Briefly, z-stack images of astrocytes were captured and manually segmented using the Simple Neurite Tracer plugin for ImageJ (Longair, Baker, & Armstrong, 2011; Schindelin et al., 2012). Segments were maximally projected to form a composite, segmented astrocyte, and concentric circles with increasing radii of 2 μm were overlaid and the number of intersections was counted automatically and plotted by distance by the Sholl plugin for ImageJ (Ferreira et al., 2014). Two or more tissue sections through each anatomical region were analyzed per animal (n = 6 mice per group). Sholl analysis of astrocyte morphology was conducted by persons blind to the experimental condition of the animal.

### RNA Isolation and Quantification

#### Brain tissue

Regions of interest (STR and ETC) were bilaterally microdissected (as described above) and submerged in an RNA stabilization solution (pH 5.2) at 4°C, containing in mM: 3.53 ammonium sulfate, sodium citrate, and 13.33 EDTA (ethylenediaminetetraacetic acid). Tissue was transferred to a sterile tube containing 300 μl TRI-reagent (Cat.# 11-330T, Genesee Scientific) and homogenized with a mechanical pestle before centrifuging at 13,000 x g for 3 minutes. Supernatant was removed to a new tube where 250 μl of chloroform was added and tubes vigorously shaken twice for 10 seconds followed by 3 minutes of resting on ice and centrifugation at 13,000 x g for 18 minutes at 4°C. The upper, aqueous layer was carefully removed to a new tube, an equal volume of 100% ethanol was added, and the sample was thoroughly mixed before RNA purification using the Zymo Direct-zol RNA Miniprep (Cat.# 11-330, Genesee Scientific) according to the manufacturer’s instructions. RNA was eluted in 35 μl of DNAse/RNAse free water before spectrophotometric analysis of RNA purity and concentration.

#### Primary astrocyte cultures

Culture media was removed, and astrocytes were washed twice with ice-cold DPBS before direct addition of TRI-reagent to each well (250 μl or 500 μl of TRI-reagent for 12- or 6-well plates, respectively). Cells were gently scraped with individual cell scrapers before removing to respective, sterile tubes on ice. 250 μl of chloroform was added and tubes vigorously shaken twice for 10 seconds followed by 3 minutes of resting on ice and centrifugation at 13,000 x g for 18 minutes at 4°C. The upper, aqueous layer was carefully removed to a new tube, an equal volume of 100% ethanol was added, and the sample was thoroughly mixed before RNA purification using the Zymo Direct-zol RNA Miniprep according to the manufacturer’s instructions. RNA was eluted in 35 μl of DNAse/RNAse free water before spectrophotometric analysis of RNA purity and concentration.

### Complementary DNA Synthesis and Quantitative RT-PCR

Complementary DNA (cDNA) was synthesized from either 200 ng (astrocyte cultures) or 400 ng (brain tissue) of isolated RNA using the qPCRBIO cDNA Synthesis Kit (Cat.# PB30.11-10, PCR Biosystems, Wayne, PA) following manufacturer’s guidelines before being diluted 1:5 in DNAse/RNAse free water and stored at −20°C. Gene expression changes were measured with quantitative RT-PCR (qRT-PCR) similarly as previously described (Halliday et al., 2019; Lundquist et al., 2019). Briefly, qRT-PCR was run with 2 μl of cDNA and qPCRBIO SyGreen master mix (Cat.# PB20.11-01, PCR Biosystems) on an Eppendorf Mastercycler Ep Realplex (Eppendorf, Hauppauge, NY) using a program of 15 min at 95°C, followed by 40 cycles of 15 seconds at 94°C, 30 seconds at 55°C, and 30 seconds at 72°C. Data was collected and normalized on Eppendorf Realplex ep software. Standard delta-CT analysis (Livak & Schmittgen, 2001) was used to quantify fold changes in gene expression in experimental groups normalized to controls, with Actb serving as a housekeeping gene. A complete list of primer pairs can be found in Table 2.

**Table 1.** List of antibodies used during immunohistochemical analysis.

| Antibody | RRID | Host Species | Immunogen | Manufacturer | Dilution |
| --- | --- | --- | --- | --- | --- |
| Rabbit anti-GFAP | AB_10013382 | Rabbit | GFAP isolated from cow spinal cord | Agilent (Cat# Z0334) | 1:2000 |
| Rabbit anti-SOX9 | AB_2239761 | Rabbit | KLH-conjugated linear peptide corresponding to the C-terminal sequence of human Sox9 | Millipore (Cat# AB5535) | 1:2000 |
| Mouse anti-PSD-95 | AB_2092543 | Mouse | Synthetic Peptide | Millipore (Cat# AB9708) | 1:2000 |
| Rat anti-GFAP | AB_211868 | Rat | Bovine GFAP | Millipore (Cat# 345860-100UG) | 1:2000 |
| Rabbit anti-synaptophysin | AB_2286949 | Rabbit | Synthetic peptide within Human Synaptophysin aa 250-350 (C terminal). | Abcam (Cat# ab32127) | 1:2000 |
| Mouse anti-ALDH1L1 | AB_10671695 | Mouse | Protein amino acids 1-902 (full-length) of rat Aldh1L1 | UC Davis/NIH NeuroMab Facility (Cat# 75-164) | 1:500 |
| Goat anti-mouse (488 nm) | AB_2534069 | Goat | Gamma Immunoglobins Heavy and Light Chains | Thermo Fisher Scientific (Cat# A-11001) | 1:5000 |
| Goat anti-rabbit (568nm) | AB_143157 | Goat | Gamma Immunoglobins, Heavy and Light chains | Thermo Fisher Scientific (Cat# A11011) | 1:5000 |
| Goat anti-rat (568 nm) | AB_2534121 | Goat | Gamma Immunoglobins Heavy and Light Chains | Thermo Fisher Scientific (Cat# A-11077) | 1:5000 |

**Table 2.**
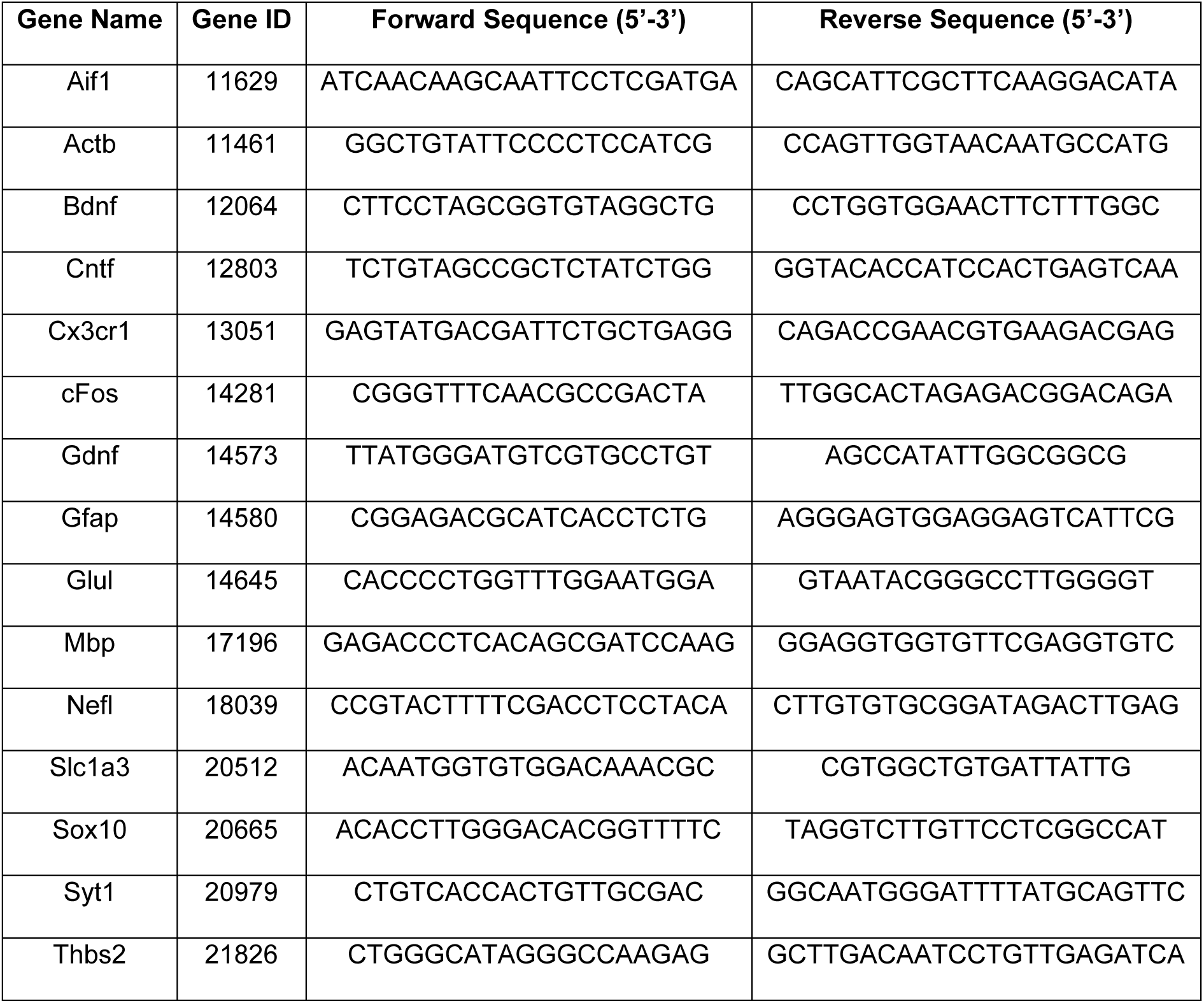
List of primers used for qRT-PCR experiments.

### Statistical Analysis

All statistical tests were carried out and graphs made in Prism 8.3 (GraphPad, San Diego, CA) with statistical significance set at p < 0.05. No sample size calculations were performed prior to the start of the study but are based on previous publications from our lab (Halliday et al., 2019; Lundquist et al., 2019). All data included was normally distributed as assessed by Shapiro-Wilk normality testing, and no data points were excluded from analysis. Simple linear regression of accelerating rotarod behavior was used to assess differences in initial coordination and learning rate between groups. Unpaired, two-tailed T-tests were used for all qRT-PCR analyses, total astrocyte process intersections in Sholl analysis, total synaptophysin- or PSD95-positive puncta, and initial coordination and learning rates on the accelerating rotarod task. Two-way ANOVA with Bonferroni’s multiple comparisons were used for analysis of Sholl plots and learning curves for accelerating rotarod behavioral analysis. For Sholl plots, distance from soma served as the within-subject comparison, and for rotarod learning curves, trial number served as the within-subject comparison. For both Sholl plots and rotarod analysis, L-lactate administration served as the between-subject comparison. Where feasible, p values are listed on the corresponding figures and all p values are listed in the results section.

## Results

### L-lactate and 3,5-DHBA administration elevates the expression of astrocyte-specific gene transcripts in primary astrocyte cultures

The effect of L-lactate administration on astrocyte-specific gene transcripts was performed in primary astrocytes cultures. Figure 2 shows the near confluence of astrocytes based on SOX9 immunohistochemical staining, demonstrating a highly enriched astrocyte culture. Gene transcript expression was assessed following an acute (1 hour, 10mM), in vitro exposure of L-lactate (see Figure 3). Following L-lactate exposure, there was an increase in the expression of *Gfap* (a marker of astrocytic reactivity; 1.49-fold increase, p = 0.0027), *Thbs2* (a glycoprotein involved in synaptogenesis; 1.84-fold increase, p = 0.027), and genes for the neurotrophic factors *Gdnf* (glial cell line-derived neurotrophic factor; 2.27-fold increase, p = 0.015), *Bdnf* (brain-derived neurotrophic factor; 1.64-fold increase, p = 0.054), and *Cntf* (ciliary neurotrophic factor; 3.22-fold increase, p < 0.001). Additionally, there was a significant increased expression of the immediate early gene *cFos* (3.71-fold increase, p = 0.0017). No significant change in expression was seen for genes involved in glutamate neurotransmission including glutamine synthetase (*Glul*) and the glutamate transporter (*Slc1a3*) (Figure 3A).

**Figure 3.**
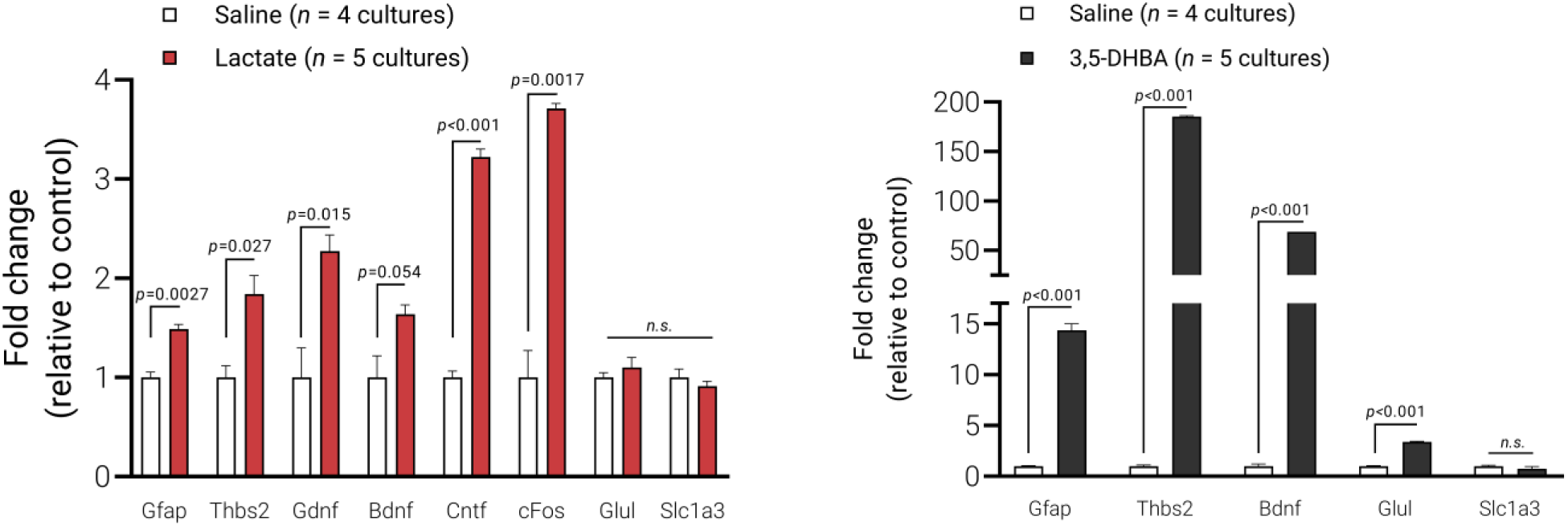
L-lactate and 3,5-DHBA administration to primary astrocytes induces changes in astrocyte plasticity gene expression. (A) Administration of 10mM L-lactate to primary astrocytes increased gene expression in several genes, as measured by qRT-PCR (*n* = 4-5 samples from five independent culture preparations). (B) Administration of 2.5mM 3,5-DHBA to primary astrocytes increases gene expression as measured by qRT-PCR (*n* = 4-5 samples from five independent culture preparations).

Following exposure to the L-lactate receptor HCAR1 agonist 3,5-DHBA (1 hour, 2.5mM), to primary astrocyte culture, there was a significant increase in the expression of *Gfap* (14.35-fold increase, p < 0.001), *Thbs2* (185.47-fold increase, p < 0.001), *Bdnf* (68.67-fold increase, p < 0.001), and *Glul* (3.39-fold increase, p < 0.001) (Figure 3B). There was no significant effect of 3,5-DHBA exposure on the expression of the glutamate transporter (*Slc1a3*).

### L-lactate administration elevates the expression of astrocyte-specific gene transcripts in normal healthy mice

L-lactate (2g/kg, estimated final concentration of 10mM) (Morland et al., 2017) or saline was administered by I.P. injection in normal mice for 10 days and astrocyte-specific gene expression was determined by qRT-PCR in the striatum (STR) and ectorhinal cortex (ETC). These two regions of the brain were selected based on the report of exercise-induced increase (STR) or no change (ETC) in regional cerebral blood flow (Wang et al., 2013). In the STR, L-lactate significantly increased the expression of genes including *Gfap* (a marker of astrocytic reactivity; 3.57-fold increase, p < 0.001), *Thbs2* (a glycoprotein involved in synaptogenesis; 6.67-fold increase, p < 0.001), and the neurotrophic factor genes *Gdnf* (15.3-fold increase), Bdnf (30.1-fold increase) and *Cntf* (82-fold increase; all p < 0.001) relative to saline treated animals. Additionally, there was a significantly increased expression of the immediate early gene *cFos* (30.1-fold increase, p < 0.001). The genes involved in glutamate neurotransmission were also increased in expression including *Glul* (glutamine synthetase; 6.44-fold increase) and *Slc1a3* (glutamate transporter; 4.38-fold increase; both p < 0.001) relative to saline treated animals (Figure 4A). Within the ETC, L-lactate administration did not result in any significant changes in gene expression compared to saline administration (Figure 4B).

**Figure 4.**
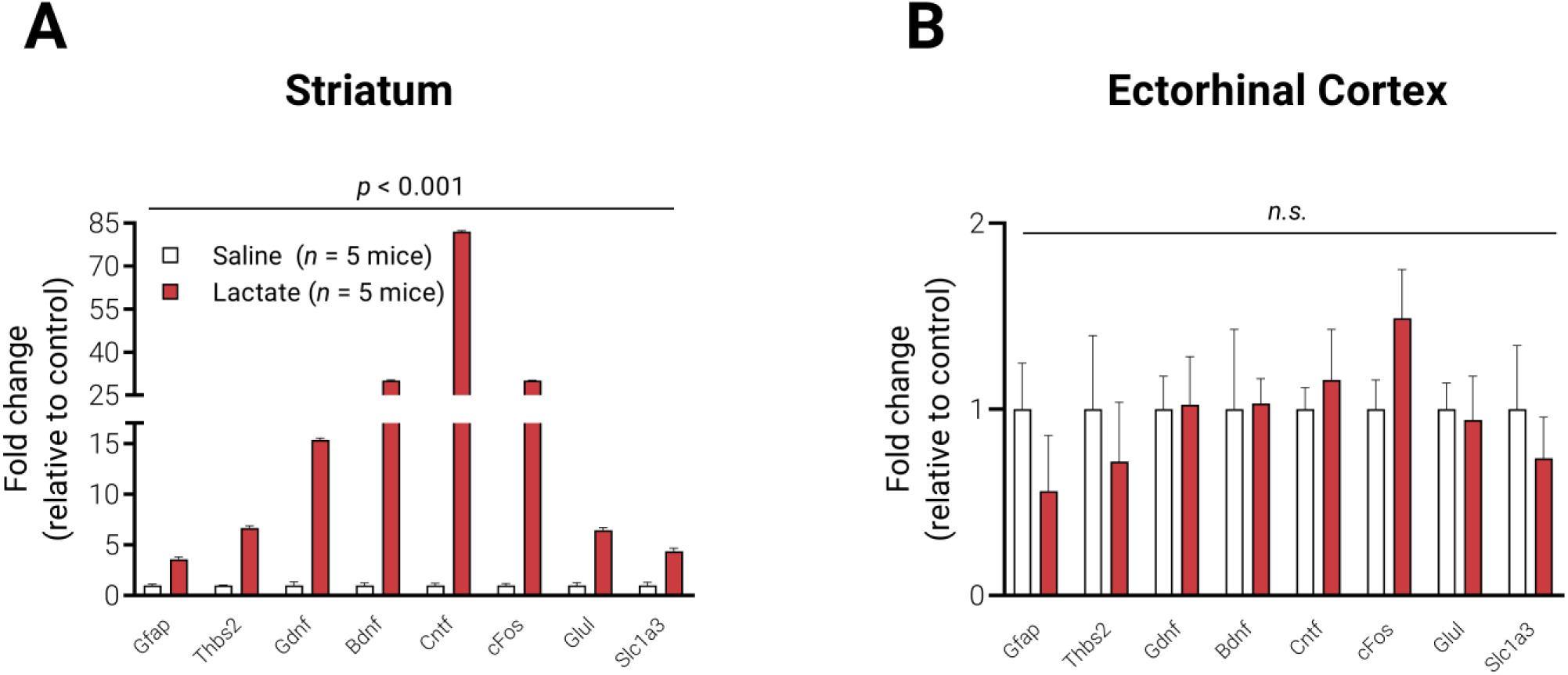
L-lactate administration induces changes in astrocyte plasticity gene expression in mice. (A) 10 days of 10mM L-lactate administration significantly increased expression of astrocyte plasticity genes in the striatum of mice, as assessed by qRT-PCR (*n* = 5 mice per group). (B) 10 days of 10mM L-lactate administration did not affect expression of plasticity-related genes in the ectorhinal cortex of mice, as assessed by qRT-PCR. *N* = 5 mice per group.

### L-lactate administration results in altered astrocyte morphology in normal healthy mice

L-lactate (2g/kg, estimated final concentration of 10mM) or saline was administered by I.P. injection in normal mice for 10 days and Sholl analysis was used to examine two parameters, including (i) astrocytic arborization and (ii) the total number of intersections in both the STR and ETC. L-lactate administration resulted in a significant increase in the arborization of astrocytic processes (n = 17 cells per group, two-way ANOVA, F(1, 1312) = 99.23, p < 0.001) compared to saline controls (Figure 5A). Similarly, the total number of intersections per striatal astrocyte was also significantly increased following L-lactate administration when compared to saline controls (n = 17 cells per group, unpaired two-tailed t-test, t = 2.882, p = 0.0070) (Figure 5B). In the ETC, L-lactate administration did not have a significant effect on the overall morphological complexity of astrocytes (n = 12-13 cells per group, two-way ANOVA, F(1, 942) = 1.674, p = 0.196) (Figure 5C) or on the total number of intersections per astrocyte (n = 12-13 cells per group, unpaired two-tailed t-test, t = 0.3768, p = 0.710) compared to saline controls (Figure 5D).

**Figure 5.**
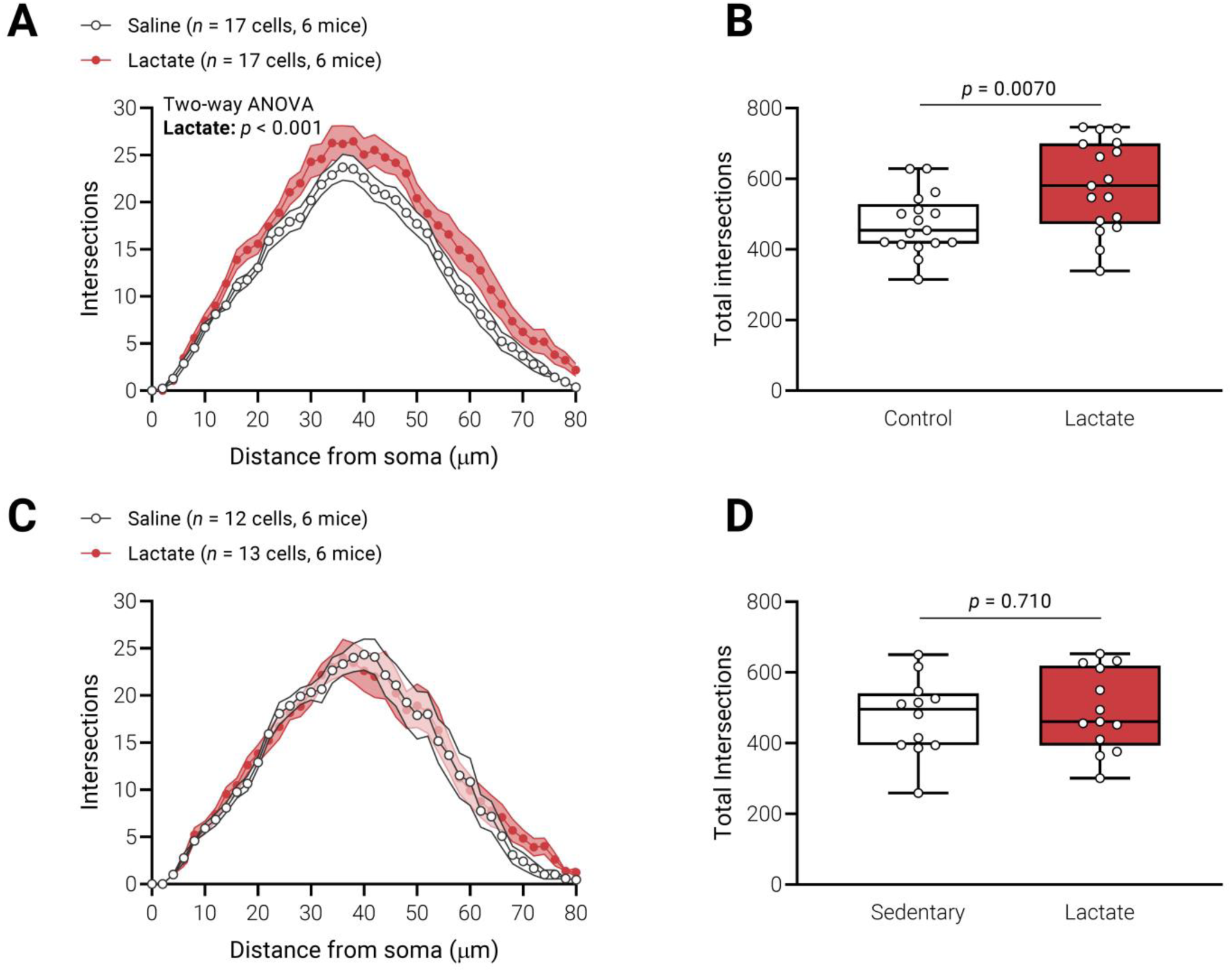
L-lactate administration causes morphological remodeling of striatal astrocytes, but not cortical astrocytes in mice. (A) 10 days of 10mM L-lactate administration in vivo significantly increased the complexity of striatal astrocyte morphology as measured by Sholl’s analysis. (B) The total number of intersections per astrocyte, as measured by Sholl’s analysis, was significantly increased in striatal astrocytes following L-lactate administration in vivo. (C) 10 days of 10mM L-lactate administration in vivo did not affect ectorhinal cortical astrocyte morphology, as measured by Sholl’s analysis. (D) The total number of intersections per astrocyte, as measured by Sholl’s analysis, did not significantly differ in ectorhinal cortical astrocytes following L-lactate administration in vivo. *N* = 6 mice per group.

### L-lactate administration does not increase striatal PSD-95 or synaptophysin protein expression in normal healthy mice

The patterns of expression of PSD-95 and synaptophysin (two proteins involved in synaptogenesis) (Toy et al., 2014) were determined by counting the number of immunopositive puncta in the STR and ETC. L-lactate administration to mice did not result in a significant increase in the amount of PSD-95 positive puncta compared to saline controls in either the STR or the ETC (n = 15-20 sections per group, STR: unpaired two-tailed t-test, t = 1.345, p = 0.188; ETC: unpaired two-tailed t-test, t = 1.003, p = 0.324) (Figure 6A-C). Similarly, there was no significant difference in synaptophysin positive puncta in these same mice following L-lactate administration compared to saline controls in either the STR or the ETC (n = 12 sections per group, STR: unpaired two-tailed t-test, t = 0.5941, p = 0.559; ETC: unpaired two-tailed t-test, t =1.714, p = 0.100) (Figure 6D-F).

**Figure 6.**
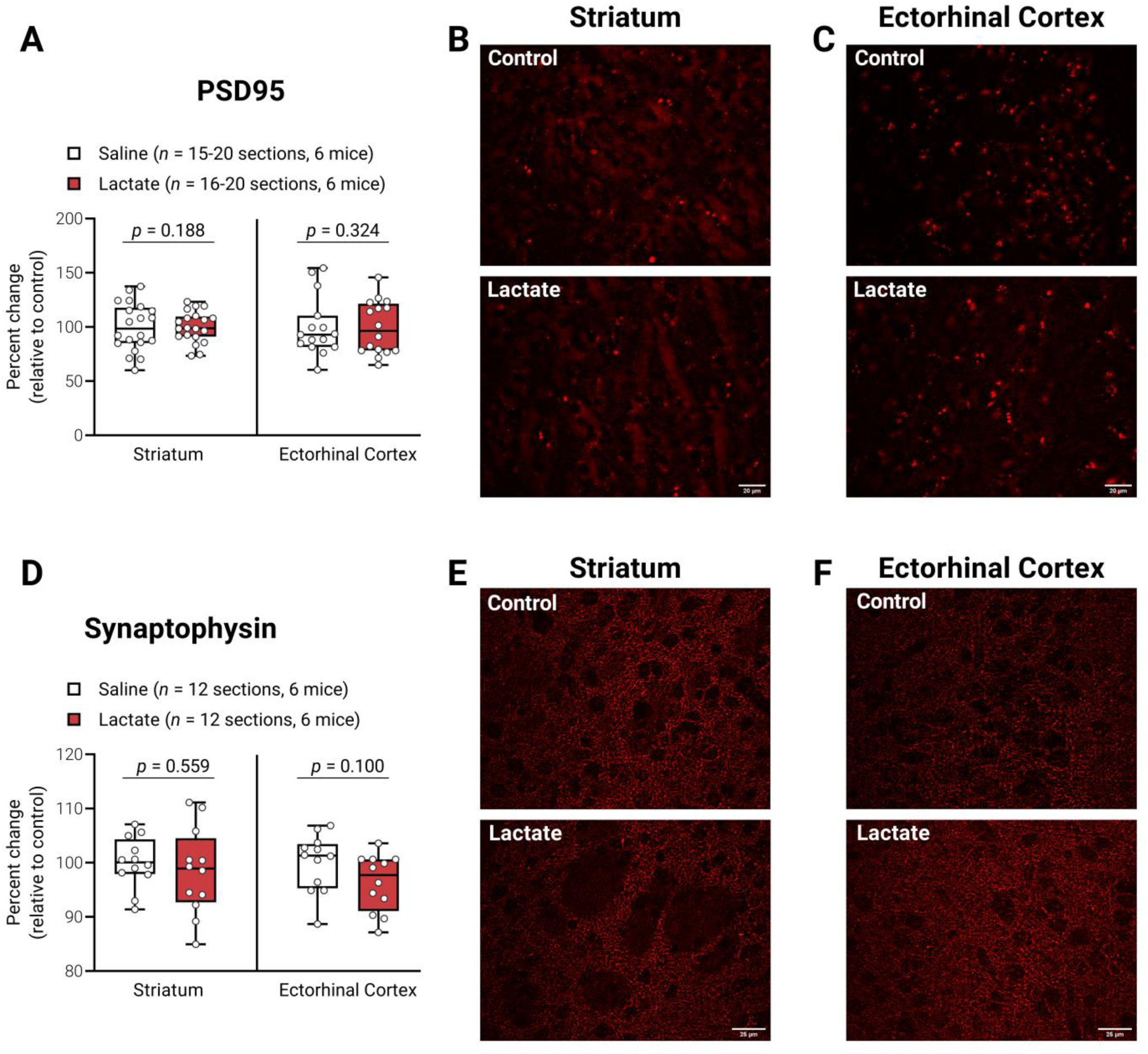
L-lactate administration does not increase synaptogenesis in striatum or ectorhinal cortex in mice. (A) PSD-95-positive puncta did not significantly differ between control and L-lactate-administered mice. (B) Representative immunohistochemistry showing PSD-95-positive puncta in the striatum. (C) Representative immunohistochemistry showing PSD-95-positive puncta in the ectorhinal cortex. (D) Synaptophysin-positive puncta did not significantly differ between control and L-lactate-administered mice. (E) Representative immunohistochemistry showing synaptophysin-positive puncta in the striatum. (F) Representative immunohistochemistry showing synaptophysin-positive puncta in the ectorhinal cortex. *N* = 6 mice per group.

### L-lactate administration does not improve motor performance on the accelerating rotarod in normal healthy mice

The effect of L-lactate administration on motor performance in mice was determined by the latency-to-fall from the accelerating rotarod. Over the 20 trial period, L-lactate administration did not significantly alter performance on the rotarod (n = 5 mice per group, two-way ANOVA, F(1, 160) = 1.490, p = 0.224) (Figure 7A-B). Additional linear regression analysis of latency-to-fall curves revealed that L-lactate did not significantly change initial coordination (unpaired two-tailed t-test, t = 1.343, p = 0.216) or the learning rate (unpaired two-tailed t-test, t = 0.992, p = 0.350) between the L-lactate and control groups (Figure 7C).

**Figure 7.**
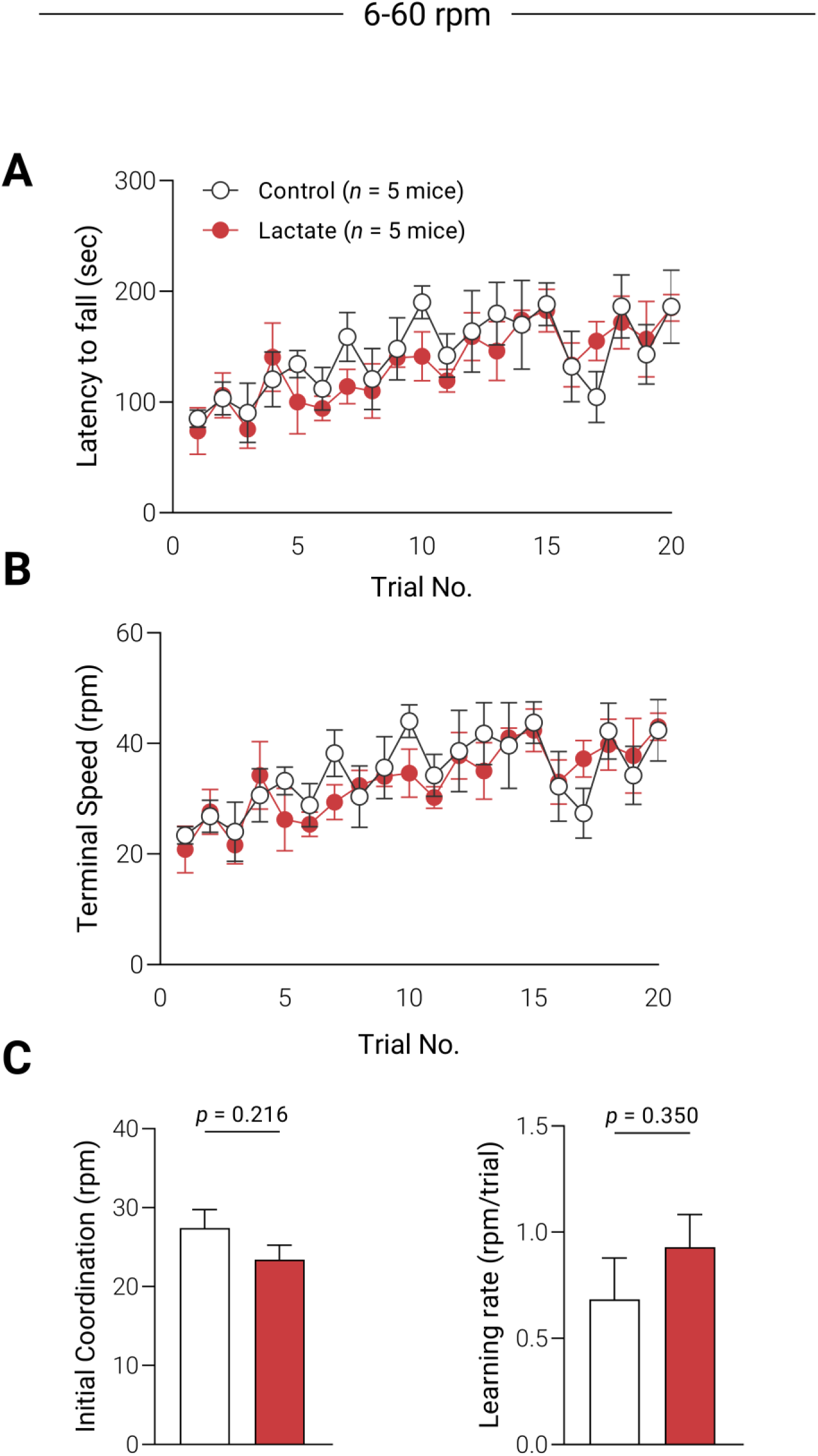
L-lactate administration does not improve motor performance on accelerating rotarod. (A) Latency to fall (in seconds) for control and L-lactate-administered mice did not differ over 20 trials of learning. (B) Terminal speed (in rpm) for control and L-lactate administered mice did not differ over 20 trials of learning. (C) Both initial coordination (in rpm) and learning rate (rpm/trial) did not significantly differ between control and L-lactate administered mice over 20 trials of learning. *N* = 5 mice per group.

## Discussion

Our studies in astrocyte cell culture demonstrated that the acute administration of a physiological dose of L-lactate increased the transcript expression of several neurotrophic factors (NTFs) including brain-derived neurotrophic factor (*Bdnf*), ciliary neurotrophic factor (*Cntf*), and glial cell line-derived neurotrophic factor (*Gdnf*). Our findings are consistent with published studies demonstrating increased level of BDNF expression in cultured human astrocytes following 4- or 24-hours of L-lactate exposure at various concentrations (5, 10, or 25mM) (Coco et al., 2013). In addition, we found that daily peripheral IP injections of 10mM L-lactate to mice significantly increased the expression of these same NTF transcripts (*Bdnf, Gdnf, Cntf*) within the STR compared to saline treated mice. Studies from others have also shown that L-lactate can lead to the increased expression of BDNF within the hippocampus, leading to increased neurogenesis and memory and learning (El Hayek et al., 2019). Our study is one of the first to report the differential effect of L-lactate on NTF expression, including *Bdnf*, within the STR. While we observed L-lactate-induced increase of NTFs in the STR there was no significant increase in expression of these NTFs in the ETC. This differential effect of NTF expression may be due to the fact that L-lactate was administered to mice during a walking task. The STR, unlike the ETC, is a brain region that subserves motor behavior that is likely activated during walking leading to increased blood flow and subsequent higher concentrations of L-Lactate. This differential effect in the STR is also consistent with studies that have similarly reported STR-specific increase in regional cerebral blood flow (rCBF) during a walking task in mice (Wang et al., 2013). Taken together, our study supports that while L-lactate may be important for inducing the expression of NTFs, the region-specific expression may be due in part to region specific neuronal activation and increased rCBF.

L-lactate acts through its G-protein coupled receptor, Hydroxycarboxylic Acid Receptor 1 (HCAR1), the primary lactate receptor expressed in astrocytes (Lauritzen et al., 2014). Our findings showed that activation of HCAR1 with the agonist 3,5-DHBA replicated our L-lactate findings by significantly increasing the expression of NTF transcripts. This finding supports that activation of the primary L-lactate receptor can mediate NTF expression via downstream signaling cascades (Liu et al., 2012; Morland et al., 2015). There are several potential mechanisms by which L-lactate could increase the expression of NTFs including the activation of transcription factors such as the immediate early gene c-Fos (Shaulian & Karin, 2002). In agreement with this hypothesis, we found that administration of L-lactate to either astrocyte cultures or to normal healthy mice increased expression of c-Fos transcripts. An alternative mechanism is through the elevated expression of immune cytokines, including TNF-alpha. Specifically, the administration of L-lactate to astrocyte cultures results in increased TNF-alpha expression (Andersson, Rönnbäck, & Hansson, 2005) and such changes in TNF-alpha can mediate increased BDNF expression (Saha, Liu, & Pahan, 2006).

Administration of L-lactate to normal healthy mice also resulted in changes in astrocyte morphology. We found that administration of L-lactate for 10 days significantly increased striatal astrocyte arborization, as defined by the number of GFAP-positive branching processes and increased the number of intersections based on Sholl analysis (Lundquist et al., 2019; Sholl, 1953). These changes in morphology were localized to the STR but not the ETC in mice subjected to walking behavior. One mechanism by which L-lactate may remodel astrocytes is through BDNF which has been shown to regulate astrocyte morphology in both the developing and adult brain (Holt et al., 2019; Ohira et al., 2007). In addition, previous studies in our laboratory have shown that exercise, in the form of treadmill running, also resulted in altered STR astrocyte morphology (Lundquist et al., 2019). It is well-established that exercise significantly elevates BDNF expression (Cotman & Berchtold, 2002; Neeper, Góauctemez-Pinilla, Choi, & Cotman, 1995). Taken together, these findings support that L-lactate-induced changes in astrocyte morphology may be mediated through BDNF.

In addition to changes in transcripts for *Bdnf* and *cFos*, we also found that L-lactate administration increased the expression of several astrocyte-specific genes including *Gfap* (a marker of astrocyte reactivity), *Thbs2* (a glycoprotein involved in synaptogenesis), *Glul* (glutamine synthetase), and *Slc1a3* (glutamate transporter EAAT1). Taken together, these genes are potentially involved in various aspects of neuroplasticity as shown in a number of models of memory and learning (Allen, 2014). For example, changes in GFAP serve as a readout for reactive astrogliosis where increases in expression correlate with several physiological characteristics including morphological remodeling (Sofroniew & Vinters, 2010). Thbs2 plays a central role in excitatory synaptogenesis through its involvement in cell-cell interactions with the postsynaptic α2-δ1 receptor (Christopherson et al., 2005; Eroglu et al., 2009). Both Glul and Slc1a3 regulate excitatory glutamatergic neurotransmission (Perego et al., 2000), manage the synaptic occupancy of glutamate (Rothstein et al., 1996), and mediate neuroplasticity as shown in models of learning and memory (Li et al., 2012).

We observed that the peripheral administration of L-lactate did not lead to changes in either synaptogenesis, as defined as changes in PSD-95 or synaptophysin, within the STR, nor improve motor behavior. This in contrast to reports demonstrating that the administration of L-lactate to the hippocampus promotes both neurogenesis and improved cognitive behavior (Lev-Vachnish et al., 2019; Rice et al., 2002).

The failure to observe improved motor behavior following L-lactate administration may be due in part to ceiling effects in normal healthy mice, whereby high regional levels of L-lactate may be needed to induce striatal plasticity. Interestingly, intensive exercise has been shown to increase central levels of L-lactate (Overgaard et al., 2012), increase STR rCBF (Wang et al., 2013) and to induce both hippocampal neurogenesis and STR synaptogenesis (Toy et al., 2014; Van Praag, Kempermann, & Gage, 1999). Therefore, the induction of STR synaptogenesis may require STR activation as precipitated by intensive exercise and the simultaneous increase in regional levels of lactate. Taken together, our findings support that peripheral L-lactate may be important in promoting the expression of astrocyte-specific genes involved in neuroplasticity (Allen & Lyons, 2018). In addition, our study suggests that L-lactate in combination with exercise, a mechanism to enhance rCBF and neuronal activation, may be needed to induce region specific change in synaptogenesis and behavioral motor change. A similar phenomenon is seen in ovariectomized rats, where the effects of estrogen replacement on BDNF expression were enhanced when combined with exercise (Berchtold, Kesslak, Pike, Adlard, & Cotman, 2001).

## Conclusion

Overall, our results are the first to show that physiologic L-lactate can initiate astrocytic-specific transcriptional and structural remodeling within the STR. However, the administration of L-lactate is not sufficient to induce mechanisms leading to synaptogenesis and improved motor behaviors. These novel findings provide a framework that links mechanisms of peripheral metabolism in muscle and neuronal activity with changes in neuronal connectivity within specific regions of the brain. Further elucidation of these mechanisms will continue to provide valuable insights into the role of astrocytes and their link to synaptogenesis. Thus, regulation of L-lactate during exercise may serve as a therapeutic modality to mediate structural, functional, and behavioral improvements to promote repair, particularly in neurodegenerative diseases such as Parkinson’s disease.

## Author contributions

All authors take full responsibility for the integrity and analysis of all data presented. *Conceptualization*, A.J.L., G.M.P., M.W.J.; *Methodology*, A.J.L., G.M.P., M.W.J.; *Investigation*, A.J.L., T.J.G., G.M.P., M.W.J.; *Formal Analysis*, A.J.L., G.M.P., M.W.J.; *Writing – Original Draft*, A.J.L., T.J.G., G.M.P., M.W.J.; *Visualization*, A.J.L., M.W.J.; *Supervision*, G.M.P., M.W.J.; *Funding Acquisition*, G.M.P., M.W.J.

## Data Accessibility

Further information regarding resources, reagents and data availability should be directed to the corresponding author and will be fulfilled upon reasonable request.

## Acknowledgments

The authors would like to acknowledge the support of the U.S. Army NETRP (Grant No. W81XWH-18-1-0666 and W81XWH-19-1-0443), the National Parkinson’s Foundation, the Confidence Foundation, the Plotkin Family Foundation, Anthony McClaren and the Climb Above Parkinson Foundation, and the Achievement Rewards for College Scientists Foundation, Los Angeles Founder Chapter. A special thanks to Friends of the USC Parkinson’s Disease Research Group including George and Mary Lou Boone, Walter and Susan Doniger, and the family of Don Gonzalez Barrera. The authors declare no competing financial interests.

